# Identifying crossovers in a cattle pangenome containing haplotype-resolved assemblies from half-siblings

**DOI:** 10.64898/2026.02.20.706955

**Authors:** Alexander S. Leonard, Hubert Pausch

## Abstract

**Background:** Recombination of parental haplotypes is a fundamental biological process that ensures proper segregation of homologous chromosomes and creates new combinations of alleles during meiosis. Crossover events are typically detected from large-scale pedigree-based genetic studies or linkage disequilibrium-based recombination maps, although these are generally limited to SNPs. Increasing amounts of long read sequencing and haplotype-resolved assemblies offer an alternative approach to examining recombination events at basepair resolution, albeit with much smaller sample sizes.

**Results:** Here, we analyse five high-quality genome assemblies from the Simmental cattle breed, including a newly assembled triobinned HiFi assembly of an Eringer x Simmental cross (N50 of 77 Mb and a k-mer quality value of 55.3). We integrate the five assemblies, of which two originate from maternal half-siblings, into a reference-free Simmental-specific pangenome. By considering path similarities in the pangenome, we were able to identify putative crossover events in the haplotypes of the half-siblings, as well as a greater number of events relative to the cousin due to an additional degree of generational separation. We validated the pangenome approach with phased SNPs called from linear alignments of maternal short read sequencing, with 23 of 30 chromosomes having the same recombination predictions. We identified 5 and 16.7 Mb of non-reference insertion sequences respectively shared or private to the half-siblings, enabling testing for recombination events beyond only SNP markers. We also identified four differentially methylated CpG clusters from the 5mC signal of HiFi reads which allowed us to narrow the window containing the putative recombination event from 35 to 20 Mb within the longest run of homozygosity.

**Conclusion:** Structural variants and methylation information identified from long read sequencing and genome assemblies may help identify recombination events in regions beyond those typically called from SNPs. Furthermore, while existing long read-based methylation calls can be noisy and report unrealistic intermediate methylation levels, 5mC methylation appears to be a promising avenue for distinguishing haplotypes in the absence of genomic variation.

## Background

Recombination is a crucial driver of genomic diversity, allowing the random exchange of genetic segments between homologous chromosomes which creates new haplotype and allele combinations. This process can improve selection of advantageous combinations and purge deleterious alleles [1]. Recombination rates vary substantially along the chromosomes [2], where recombination hotspots are short genomic regions where genetic material is exchanged at much higher rates than elsewhere. These recombination hotspots enable rapid evolution of genes, particularly those related to immunity [3]; oppositely, some regions may be repressed for recombination, like supergenes [4]. A detailed understanding of recombination maps can improve estimates of variant linkage disequilibrium used in imputation and other downstream genomic analyses [5].

Recombination rates are commonly evaluated in cattle using sparse, SNP-based genotype arrays in large-scale pedigreed mapping cohorts [2, 6–8] or linkage disequilibrium [9] approaches. While these methods often account for effective population size and demographic history, their resolution is limited by the density of markers and are often lacking complex or repetitive genomic regions. Recombination events can also be detected directly from short read sequencing [10] or Hi-C reads [11, 12] from single gametes.

Recombination is also a source of structural variation, particularly through nonallelic homologous recombination. These rearrangements can lead to phenotypic disorders [13], although they can be challenging to interrogate with short read sequencing as they often fall in repetitive regions. Long reads and genome assemblies have improved the resolution of complex recombination events such as recurrent inversions [14] and transposable elements [15]. Pangenomes can be used to evaluate regions of similarity and differences across genome assemblies without the explicit need for a reference [16]. Haplotype-resolved cattle assemblies have previously been integrated in pangenomes, but primarily for assessing gene presence/absence [17, 18] or putatively trait-associated structural variants [19, 20]. Recent advances in pangenome alignment account for recombination [21, 22], utilising haplotype combinations that are not explicitly present in the graph.

Variation in methylation, primarily the 5mC marker, has also been associated with mediating or limiting recombination in arabidopsis [23], barley [24], and cannabis [25]. Although there is not a universal consensus, some studies report transgenerational inheritance of methylation patterns in mouse [9] and grape [26], as well as in human sperm for another methylation marker m6A [27]. Methylation is also an increasingly promising source of haplotype-specific information, allowing improved phasing of sequencing reads or variants [28]. However, methylation information has not yet been used to investigate recombination events in cattle.

Here, we investigate crossover events in five highly contiguous and near-complete Simmental genome assemblies. While the number of assemblies is far too small to confidently identify recombination hot- or coldspots, we demonstrate the possibility of identifying crossover events with structural variants or methylation. These structural variants and methylation markers, readily accessible from recent long read sequencing, may help identify crossover events within SNP-based runs of homozygosity.

## Methods

### Genome assemblies

We used 4 publicly available Simmental assemblies (Table 1) and assembled an additional genome from a Eringer x Simmental cross. DNA was prepared from blood samples of the F1 and cryopreserved semen samples of the Eringer sire. We generated 101 Gb of HiFi reads with a read N50 of 16.4 Kb from one PacBio Revio 25M SMRT cell. We also generated 83 and 62 Gb of Illumina 2×150bp paired-end reads for the F1 and sire respectively, with 70 Gb dam short reads publicly available (SAMEA115121771). We followed the assembly and quality assessment approach described in [18]. Briefly, we assembled HiFi sequencing reads with hifiasm [29] v.0.19.9 with parental k-mers from short read sequencing to produce haplotype-resolved assemblies. We scaffolded the contigs to the cattle reference genome, ARS-UCD2.0, and assessed gene completeness with compleasm [30] v0.2.6 and base-level correctness with merqury [31] v1.3.

**Table 1.** Simmental genome assemblies used in pangenome construction. The USA_1 assembly is used as the Simmental reference genome. NG50 is calculated with respect to a 3 Gb genome size.

| Assembly | Related | Accession | Read technology | Phasing | NG50 |
| --- | --- | --- | --- | --- | --- |
| Halfsib_1 | Yes | PRJEB42335 | HiFi | Haplotype-resolved | 46.9 |
| Halfsib_2 | Yes | this study | HiFi | Haplotype-resolved | 76.7 |
| Cousin_1 | Yes | PRJEB42335 | HiFi | Haplotype-resolved | 84.3 |
| Swiss_1 | Yes | PRJEB72196 | CLR | Primary | 15.4 |
| USA_1 | No | PRJNA677947 | ONT | Haplotype-resolved | 70.6 |

We determined whole-genome pairwise similarity using mash [32] v2.3, with 10k sketches of k-mer length 31. We then created a tree using the UPGMA linkage algorithm from SciPy [33] v1.15.2.

### Pangenome construction

We constructed per-chromosome pangenomes with pggb [16], using wfmash commit 9c63d98, seqwish commit 0eb6468, smoothxg commit e93c623, and GFAffix v.0.2.0, with a segment length of 5000 and an identity target of 0.975.

### Jaccard distance-based crossover identification

We assessed binned Jaccard distances across the pangenome as described in [19]. Briefly, we binned the “USA_1” Simmental haplotype into 1 Kb bins and extracted pangenomic subgraphs corresponding to each bin using the extract command from odgi [34] v0.9. We then calculated pairwise Jaccard distances on each subgraph using odgi similarity.

We created the Jaccard distance tree using the same UPGMA approach as for the mash distances, instead using the median Jaccard distance over the whole genome for the input matrix.

### SNP crossover identification

We aligned short sequencing reads from the Simmental dam (SAMEA115121771) of the half-sibling F1s against the “USA_1” Simmental haplotype as the reference using Strobealign (v0.15) [35]. We then called SNPs with DeepVariant [36] v1.8, using the “WGS” small model. We used BCFtools (v1.21) [37] to filter for biallelic SNPs and subset variants.

To derive variants for both half-siblings and the cousin assembly, we used vg deconstruct (v1.63.0) [38] on each chromosome pangenome, followed by vcfwave (v1.0.12) [39] to simplify variant representation.

### Pangenome alignment of short reads

To allow feasible mapping, individual chromosome graphs were simplified by only retaining nodes relevant to the reference path or the two half-sibling assemblies, followed by removing non-reference stubs with “vg clip -s”. We further simplified the graphs with “vg simplify -i 0 -L 0.8 -k”. We then merged all chromosomes (autosomes and X) together using vg ids and vg combine to produce a single whole-genome GFA.

We used vg autoindex with “-w sr-giraffe” and vg giraffe [40] in GAF mode with “--named-coordinates” to align the Simmental dam short sequencing reads previously used in the SNP analysis. We used vg surject to get linear-reference coordinates for the alignments for comparisons and annotating pangenome plots made with BandageNG (v2025.5) [41].

### Methylation crossover identification

We aligned HiFi sequencing reads for both half-sibling F1s using pbmm2 (v1.17.0) against the “USA_1” Simmental haplotype as the reference. Reads were phased using triocanu (v2.3) using k-mers from short read sequencing of their respective parents. We assessed per-CpG methylation only for the maternal assigned reads using pb-CpG-tools (v3.0.0) using a minimum coverage of 10 reads. We then took the absolute difference in CpG methylation at sites present in both F1s to assess methylation similarity. We used MethBat (v0.14.2) joint-segment to identify regions of allele specific methylation.

## Results

We collected one primary and three haplotype-resolved Simmental genome assemblies from four publicly available samples. An additional HiFi-based Simmental haplotype-resolved assembly was generated in this study from an Eringer x Simmental crossbred animal (**Table 1**). Briefly, this assembly was extremely high quality (**Supplementary Table 1**), with an NG50 of 77 Mb, an estimated gene completeness of 99.64%, and an estimated base-level correctness of 99.9997% (QV∼55.3). Three of the haplotyped-resolved assemblies were from closely related cattle, with two half-siblings sharing the same dam and the other as their first cousin (sharing a grandsire). The other assembly from a Swiss Simmental individual was distantly related to the three closely related ones, although separated by many generations. The remaining assembly from an American Simmental cow had no discernible pedigree connection with the other animals (**Figure 1a**). We confirmed the expected relationship of the animals with a mash tree based on k-mers in the genome assemblies (**Figure 1b**), with the half-siblings and cousin clustering together.

**Figure 1.**
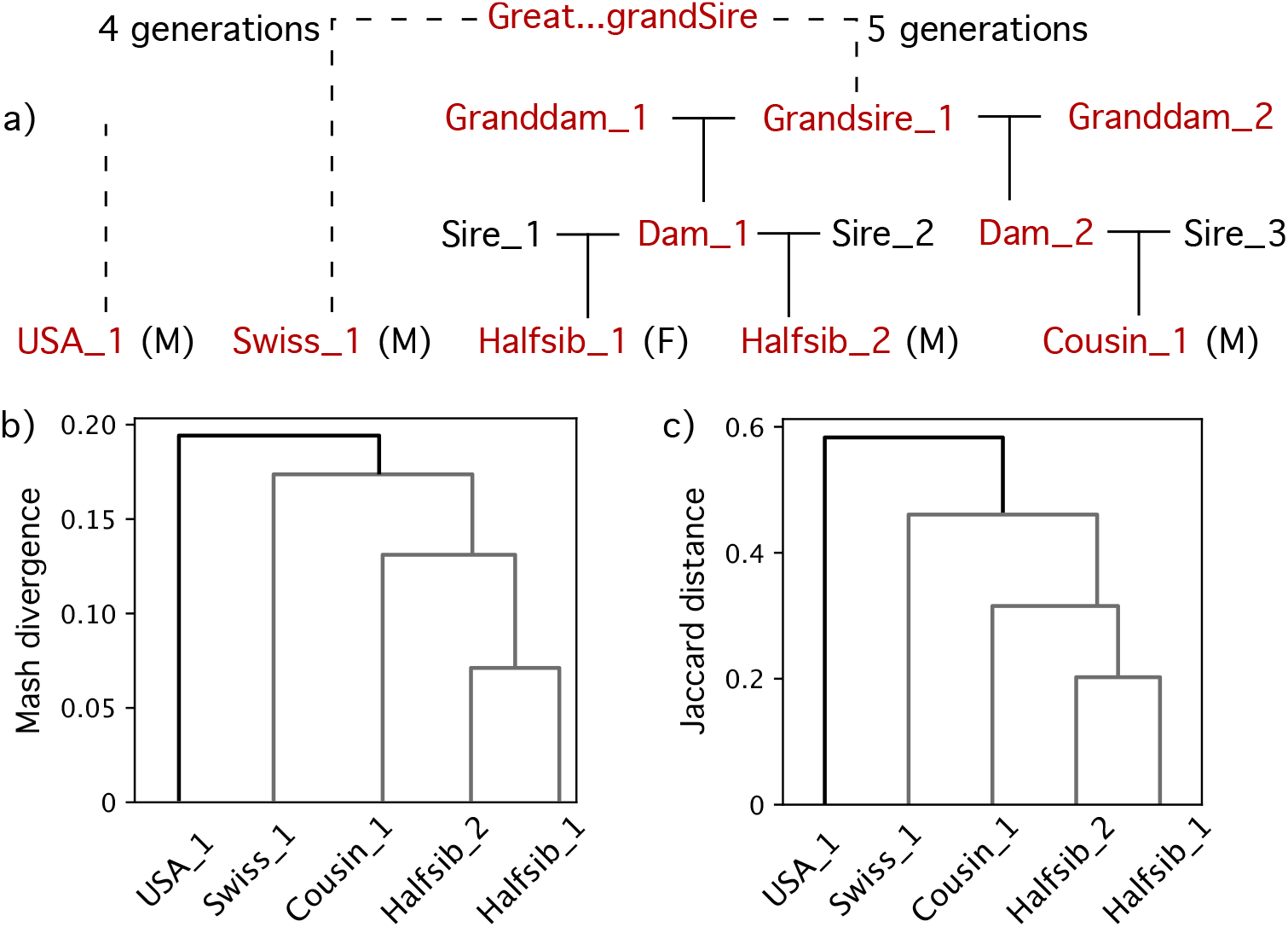
(a) Pedigree of samples derived from herdbooks where available. Simmental animals are highlighted in red and the gender of the F1s is indicated as male (M) or female (F). Swiss_1 is distantly related to the F1s through several generations of separation, while USA_1 has no presumed relationship to the others, and so their pedigrees are indicated with a dashed line. (b) Relationship tree based on the pairwise mash distance of assemblies. The cluster corresponding to the known related animals is coloured in grey. (c) Relationship tree based on the pangenome pairwise Jaccard distances of assemblies.

### Detecting recombination points in a pangenome

We used pggb to construct per-chromosome pangenomes across the 29 autosomes and X chromosome. Coincidentally, the four haplotype-resolved Simmental assemblies were all the maternal haplotype of three male F1s and one female F1, and so we excluded the Y chromosome from consideration. Despite the graphs only containing five assemblies of the same breed, we identified 443 Mb of “non-reference” sequence in total with respect to the “USA_1” reference haplotype (graph length of 3.09 Gb and reference length of 2.64 Gb). As previously reported [19], we confirmed this is largely due to unaligned centromere “tips” in the graph resulting from cattle chromosomes being acrocentric, with 259 Mb of the non-reference sequence contained in 60 massive nodes. The graphs otherwise contained 27.4 M nodes and 37.2 M edges (for an average degree of 1.37), comparable to other bovine graphs generated by pggb [43].

We quantified genomic similarity between the five Simmental assemblies using the pairwise Jaccard distance calculated over the per-assembly paths the graph for 1 Kb windows (with respect to the “USA_1” reference). The distance is close to 0 when two assemblies take the same paths throughout the chosen range and approaches 1 if they share no nodes (weighted by the length of the nodes). As such, we were able to construct another tree based on pairwise distances (**Figure 1c**), recovering the topology of the previous k-mer-based tree. However, the distance-based tree suggested a substantial (but consistent) higher genomic divergence, roughly three times the estimated mash divergence. This potentially resulted from poorly constructed or complex tangles in the graph, leading to low node similarity despite sharing similar sequence, or structural variants larger than 1 Kb. There were (11.4±3.0)×10^3^ of such 1 Kb bins with a Jaccard distance above 0.9 (approximately 11 Mb or 0.4% of the reference genome length), indicating such regions are comparatively rare.

The haplotype assemblies constructed from the half-siblings were nearly identical to each other over large stretches of each chromosome but typically had at least one distinct break where the subsequent Jaccard distance was closer to that expected for unrelated assemblies (**Figure 2a, Supplementary Figure 1**). These breaks likely correspond to recombination events, where the “maternal haplotype” assembled for each of the half-siblings corresponds to a different haplotype of the diploid maternal genome, changing from identical by descent (IBD) to non-IBD. We did not observe such a pattern in the unrelated assemblies, supporting that this method is sensitive only to real regions of IBD. While we averaged the Jaccard distance over 1 Kb windows to help mitigate small assembly errors, the true recombination resolution of the pangenome is limited only by the presence of heterozygous sequences. We re-examined two putative recombination events on BTA9 shown in Figure 2a, confirming the distinct change in heterozygous sequence between the half-siblings (**Figure 2b**) with the presence or absence of half-sibling-specific nodes. In one of those events, a 1.1 Kb heterozygous SV was within 7 Kb of the putative recombination point, further validating the heterozygous SNPs nearby and demonstrating SVs can also tag haplotype-of-origin.

**Figure 2.**
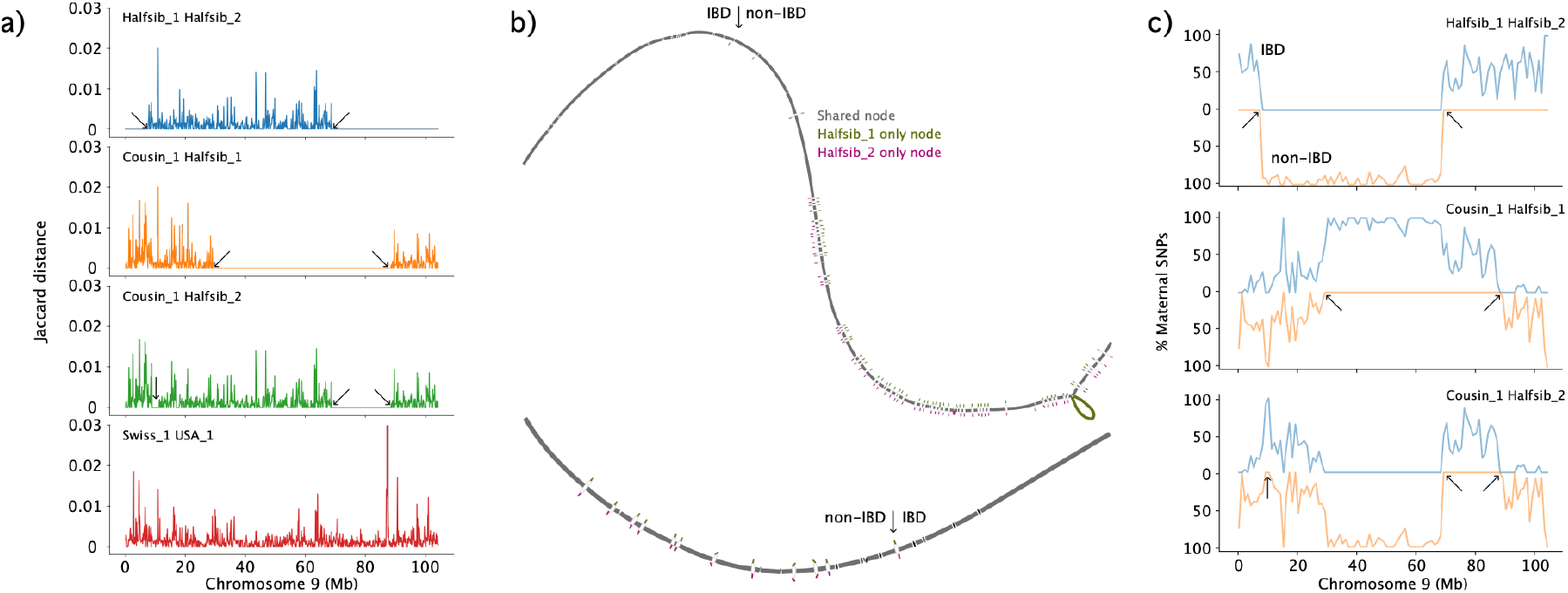
(a) Path Jaccard distance binned every 100 Kb across four pairs of assemblies on BTA9. Long stretches at zero indicate near-identical paths, with putative recombination events marked by arrows. The Swiss_1 & USA_1 pair have no expected relationship, hence have non-zero values along the entire chromosome. (b) Bandage plot for BTA9 subgraphs for the first (top) and second (bottom) recombination events between the half-siblings. Arrows mark the node-level resolution of recombination events, as determined by the presence of half-sibling specific nodes. (c) Heterozygous maternal SNPs binned every 100 Kb where the two assemblies had the same (blue) or opposite (orange) genotype. Arrows mark the same coordinates as predicted from recombination events in (a).

Based on distinct breaks in Jaccard distance (e.g. several consecutive bins are zero followed by nonzero bins) between the assemblies of the half-siblings, we estimated a mean of 1.9 putative crossover events per chromosome (**Supplementary Table 2**), close to the expectation of 2 meiotic crossovers for half-siblings (one crossover event for each dam-offspring pair). Crossovers were less frequent in shorter chromosomes, although we observed several longer chromosomes (10 and 12, respectively 102.5 and 88.5 Mb) without any putative recombination events. Detecting crossovers on the X chromosome was challenging due to several regions of suboptimal graph construction in complex regions. Likewise, the Cousin_1 assembly had multiple smaller stretches of near identical and average similarity compared to the haplotypes of the half-siblings, corresponding to multiple putative recombination events (2.9 and 2.8 events per chromosome respectively). The larger value compared to between the half-sibling assemblies is expected, given the additional generation of separation, but is still an underestimate as we cannot detect recombination within the non-shared haplotype within the cousin.

We investigated the sensitivity of detecting crossover events with the pangenome approach by considering heterozygous SNPs called from short read sequencing of the half-siblings’ dam. Both half-sibling haplotypes overwhelmingly inherited the same allele up to the 1 Kb bin containing the putative crossover, and subsequently inherited opposite alleles (**Figure 2c Supplementary Figure 2**), validating the predicted haplotype inheritance. Even the X chromosome graph prediction was validated, demonstrating the graph approach is robust even over complex regions. Several recombination events predicted by the graph-based Jaccard distance were due to runs of homozygosity (RoHs), identified by the absence of heterozygous SNPs, and so the number of predicted crossover events per chromosome was reduced from 1.9 to 1.5 (**Supplementary Table 2**). However, predictions matched between the graph and SNP approach for 23 out of the 30 chromosomes considered. Only chromosomes 5 and 17 had predictions differing by more than two predicted recombination events, with the graph potentially overestimating 3 and 4 recombination events respectively.

Due to the additional degree of relationship separation for the cousin assembly, it was possible to observe both IBD and non-IBD heterozygous SNPs in close proximity. However, we still observed extended stretches of consistent phasing. Since X-chromosomal inheritance is different to autosomal inheritance, we observed more distinct phase switches between the cousin and the half-sibling assemblies (i.e. little mixing of homozygous alternate and heterozygous SNPs). The only exception was in the X-PAR, which displayed similar behaviour to the autosomes as expected given the less restricted recombination in this region.

We also investigated the sequence context of the putative recombination events and differences in the *PRDM9* alleles, the gene associated with recombination hotspot positioning. Using the 15 bp long consensus “NCCNCCNTNNCCNCN”-motif of human *PRDM9* allele A [44] (where N represents an unconstrained nucleotide), we identified 87,848 possible matches across the USA_1 reference genome. Out of the 92 putative recombination spots, the median distance to a binding motif was 11.1 Kb, with six spots within 1 Kb.

However, the proximity was not statistically significant, with randomly permuted hotspots showing a similar distribution of proximity to binding motifs (one-sided “less” Mann-Whitney U test p=0.154). The lack of significance might be due in part to the low specificity of the binding motif and possible human-cattle *PRDM9* allele differences. We also examined the allelic sequence of *PRDM9*, identifying the gene sequence from the annotated cattle reference genome ARS-UCD2.0 in the Simmental pangenome (**Supplementary Figure 3**). The four Swiss Simmental assemblies had identical sequence across the 560 bp span consisting of five 84 bp tandemly repeated motifs (with 81 and 59 bp partial motifs preceding and following the main repeats). The USA_1 assembly contained a single T-to-C substitution at the 81^st^ base of the 3^rd^ full tandem repeat motif.

### Pangenome alignment distinguishes maternal runs of homozygosity

When comparing the pangenome- and SNP-based crossovers, we noticed it was not possible to distinguish extended RoHs in the dam from the half-siblings inheriting the same maternal haplotype. We identified three such extended RoHs of roughly 10 Mb or longer from maternal SNPs called from short read sequencing. In regions of obvious recombination, heterozygous SNPs were substantially more common than heterozygous insertions or deletions (indels). However, in all three RoHs, we observed heterozygous indels outnumbered heterozygous SNPs (**Supplementary Table 3**). The majority of these indels proved to be putative assembly errors, typically in homopolymers or other “off-by-one” differences in long SVs that could not be manually validated (**Supplementary Figure 4**). In several instances, indels from the non-Simmental parent of the crossbred F1s were erroneously included in the Simmental haplotypes, due to limited phasing power in the local region (**Supplementary Figure 5**). Another instance of a 2.3 Kb insertion unique to the Halfsib_1 haplotype assembly was later identified in both half-sibling haplotypes, where the variant was lost from the Halfsib_2 haplotype assembly during the vcfwave postprocessing of the pangenome (**Supplementary Figure 6**).

For completeness, we investigated if using short sequencing reads from the dam aligned directly to the whole-genome pangenome, combing the 29 autosomes and X chromosome built earlier into one graph, would distinguish maternal RoHs from when the haplotypes were IBD. We were able to manually verify several SVs that were either in one or both half-sibling assemblies, corresponding to a heterozygous or homozygous SV respectively. In the heterozygous case, we observed the dam short read graph alignments taking two distinct paths through the SV bubble (**Figure 3a**), while the homozygous alternate cases only had a single consistent path (**Figure 3b**). We also observed cases where the two assemblies shared the same path, but the maternal sequencing also supported an alternative path, indicating the dam was heterozygous while both half-siblings inherited the same allele (**Figure 3c**). We generalised this approach by classifying individual alignment paths as congruent with the two half-sibling assemblies or not, enabling us to identify when maternal sequencing supported an alternative path to either assembled haplotype (**Figure 3d, Supplementary Figure 7**). We found even coverage of read-to-pangenome alignments across the chromosomes, demonstrating the above effect was not due to hard-to-align regions or other causes of alignment dropout. In combination with the path Jaccard distance, we could now classify regions as maternal RoHs, IBD, or non-IBD solely through the analysis of or alignment to the pangenome without the need for any linear reference-based approaches.

**Figure 3.**
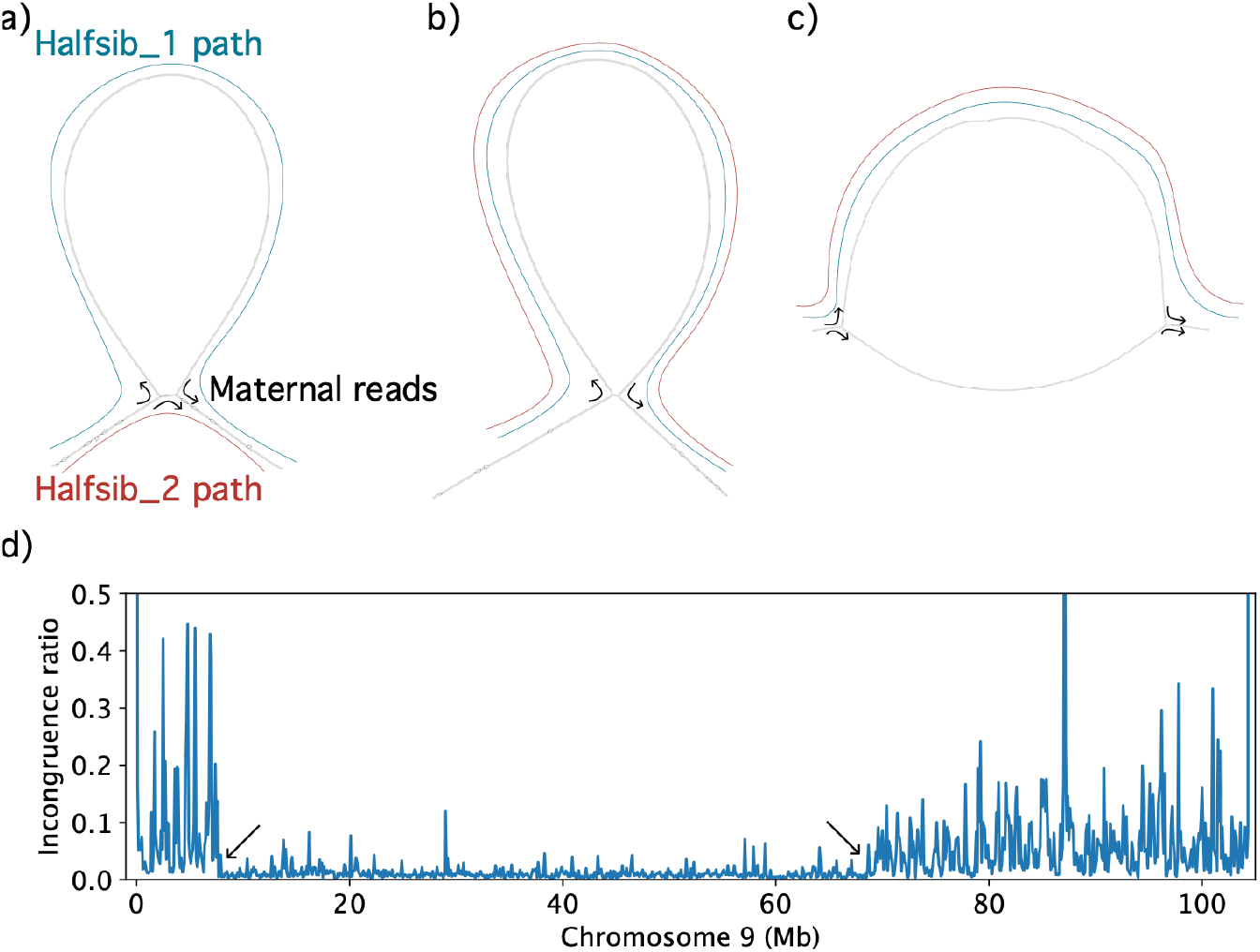
Maternal short read sequencing alignments to the graph indicate support for (a) heterozygous, (b) homozygous alternate, or (c) variants that are heterozygous in the dam but both half-siblings inherited the same allele. Blue and red paths respectively indicate the pangenome path for the Halfsib_1 and Halfsib_2 assemblies, while the black arrows indicate explicit edge junctions supported from maternal short read sequencing aligned to the graph. (d) Incongruence ratio is the ratio of maternal alignments that follow a graph path found in either half-sibling to those that don’t across 1Kb bins. A low incongruence ratio indicates that both maternal haplotypes are well represented in the graph, while a high value suggests the half-siblings are IBD and represent only one of two distinct maternal haplotypes. Arrows mark the same recombination events from Figure 2a.

### Distinguishing haplotypes with 5mC methylation

We investigated the 5-Methylcytosine (5mC) signal from the HiFi reads, hypothesising different epigenetic modifications might enable to distinguish between homozygous haplotypes. Both DNA samples used for HiFi sequencing to produce the half-sibling assemblies were extracted from blood and at similar ages, reducing possible bias in tissue-specific or age-mediated methylation. We assessed 16.4 million reference CpG sites called in both half-siblings (24.6M in Halfsib_1 and 19.3M in Halfsib_2). Of these, 12.3M were highly methylated (>70%) in both, while 1.3M where unmethylated (<30%). Overwhelmingly, methylation status was similar between both (**Supplementary Figure 8)**, with 14.9M (91% of all shared CpGs) having an absolute difference of methylation status below 30%, with only 0.2M (1.3% of all shared CpGs) sites with an absolute difference above 70% and considered to be differentially methylated sites (DMS). However, most of these sites involved mutations to the CpG motif, either removing or introducing a CpG motif rather than only modifying the epigenetic context and hence is strongly correlated with the heterozygous SNP analysis already presented (**Figure 4a**). We were unable to identify a genome-wide signal of recombination after only considering epigenetic changes to conserved CpG sites (**Figure 4b**).

**Figure 4.**
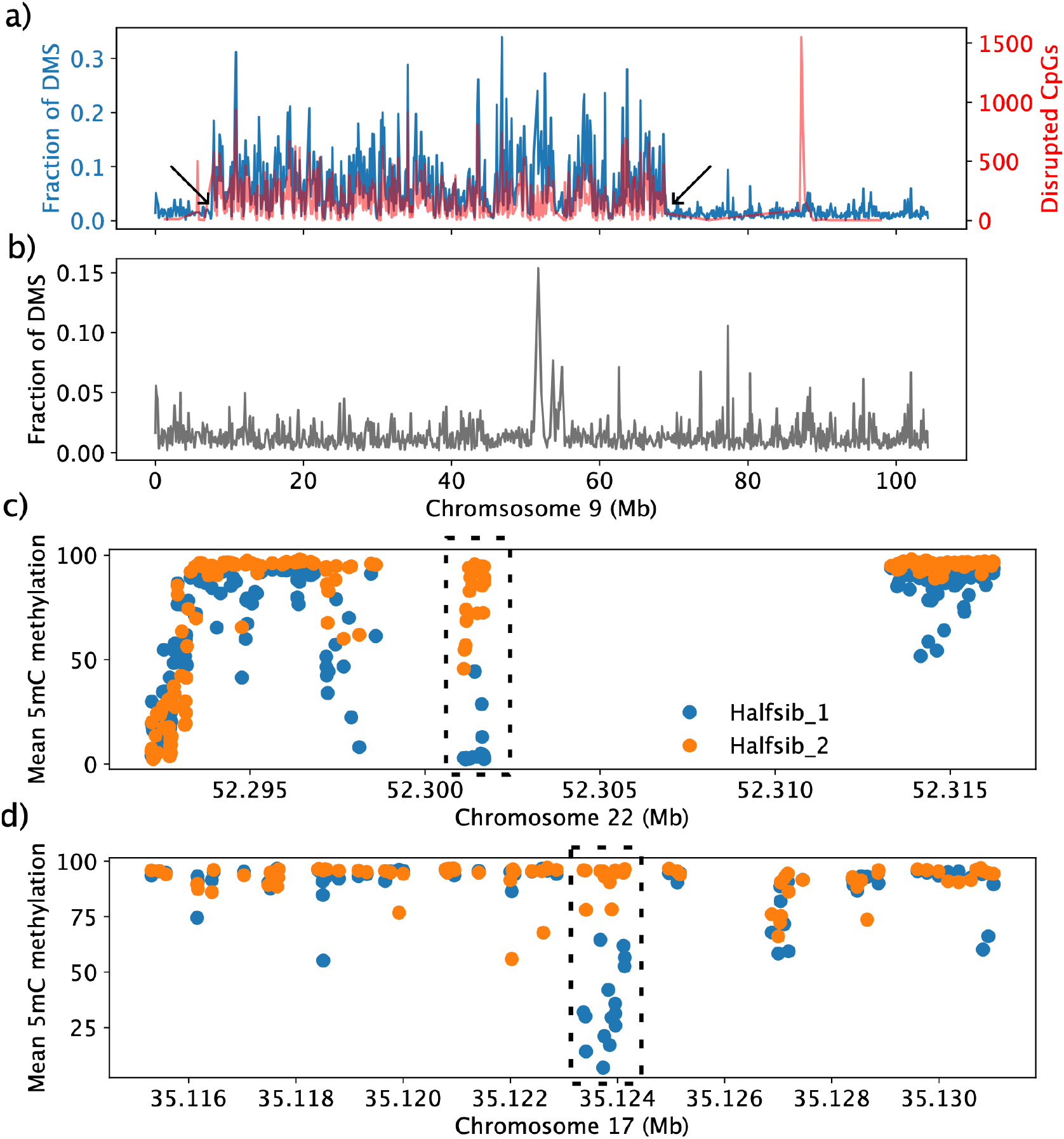
(a) The fraction of differentially methylated sites (DMS) to total CpGs in 100 Kb windows recovers the recombination events identified with pangenome similarity and SNPs, with the arrows taken from Figure 2. However, the signal is largely driven by the number of SNPs involving CpG motifs (either removing or adding), shown on an alternative y axis. (b) After excluding disrupted CpG motifs, the signal of recombination events is lost. (c-d) Mean 5mC methylation scores per CpG, averaged over all reads for each half-sibling, in a (c) expected IBD region on BTA22 and the (d) RoH on BTA17. Boxed regions indicate the detected allele-specific methylation between the two half-siblings.

We investigated genome-wide signals of allele-specific methylation (ASM), using MethBat segment on a pseudo-diplotype of the two Simmental half-siblings to identity regions of interest without limiting the analysis to a priori defined windows (i.e. CpG islands which are not defined for the USA_1 reference). There were 207 ASM windows, ranging from 102 to 3,267 bp, with a mean length of 901±611 bp. Of these, 139 (67%) occurred in non-IBD regions while 68 (33%) fell within IBD regions. Approximately 54% of the USA_1 reference genome was within non-IBD regions (1.43 out of 2.64 Gb), leading to a statistically significant result that ASM was depleted in IBD regions (One-sided Fisher’s exact test p=9.4×10^-6^). We confirmed the effect by also randomly permuting the windows 100 times (**Supplementary Figure 9**), calculating a statistically significant z-score of p=0.0039, indicating the real ASM windows occur within non-IBD regions more commonly than by chance. Despite the depletion of ASM in IBD regions, there are compelling cases of ASM within regions that are predicted to be IBD based on genetic sequence (**Figure 4c**). These may indicate non-inheritance of 5mC methylation state at some loci or biases like cell types or developmental stage affecting methylation at these loci.

Manually inspecting the alignments, we did not find evidence of a haplotype-switching event (i.e. were IBD between half-siblings before and after the RoH) in the long RoHs on BTA3 and BTA7. The BTA17 RoH (34.7 Mb long), however, changes from heterozygous to IBD between the half-siblings. This putatively suggests a crossover event occurred which was not detectable from variation in the genomic sequence, although the methylation signals can be dominated by noise. We further manually identified four candidate regions within the BTA17 RoH with clustered CpGs displaying hypermethylation in the maternal reads of one half-sibling but hypomethylation in the other (**Supplementary Table 4**). The last ASM we observed in this RoH was at 35.1 Mb, nearly 15 Mb after the start of the RoH, putatively narrowing the window where a recombination event might have occurred (**Figure 4d**). We similarly identified two additional candidate regions within the smaller BTA3 RoH (9.5 Mb long), where the half-siblings presumably remained in opposite phases. We only identified one weak candidate region within the BTA7 RoH (45.2 Mb long), as expected given the half-siblings presumably remained in the same phase. Without methylation information of the dam itself, we could not assess consistency in phase for differential methylation, instead only quantifying the magnitude of the absolute difference.

We were also able to use the 5mC methylation signal within regions containing opposite haplotypes but limited number of heterozygous SNPs to further improve phasing. We used SNPs and the long reads to phase variants into 3,430 haploblocks, located within the heterozygous regions. With pomfret, we identified 10 possible joins supported by the methylation signal in the absence of genomic variants (**Supplementary Table 5, Supplementary Figure 10**), with an average of 31±6 informative “meth-mer” sites flanking the ambiguous phase block. Given the lack of phase blocks in homozygous regions, we were unable to assess methylation-based phasing in those regions.

## Discussion

We investigated the feasibility of using genome assemblies in a pangenome for assessing recombination events. While these types of data will unlikely reach the same scale of array-genotyped cohorts, and thus be less able to predict recombination hot- or coldspots, we demonstrate that structural variants and 5mC methylation can also be used to inform recombination events. Through the reference-free pangenome approach, we also identified 21.7 Mb of insertion sequence found in either of the half-sibling assemblies, 5 Mb of which was in both haplotypes, which could contain phasing-relevant variation. However, pangenome approaches can depend on the style and quality of the graph to accurately quantify relationships between all pairs of assemblies [43], and so more robust methods and validation may be needed to separate misassemblies from true recombination events.

Although we did not observe any recombination events that could not be predicted from SNPs, potentially due to the small sample size (n=2 haplotypes), considering additional sources of genomic or epigenetic variation may help resolve large runs of homozygosity. In some species, like European Bison, over half of the genome may be in (SNP-based) runs of homozygosity but still contain comparable amounts of structural variants to other bovine species [45]. Variable number tandem repeats have a higher average mutation rate than SNPs [38] and are more likely to cause de novo mutations and interrupt RoHs, but are currently overlooked in recombination analyses. Although the assemblies examined here were also of high quality, homopolymer errors dominated putative heterozygous indels within RoHs. Incorporating sequencing reads with limited homopolymer bias for polishing or improved basecalling may be useful to reduce false positives in these regions. Similarly, the pangenome approach is sensitive enough to identify gene conversion events or other short recombination events, but the current specificity is limited by the assembly base-level accuracy.

Read-to-graph alignments could also be used with an alternative pangenome design, including both haplotypes of a sire and mapping long sequencing reads from the germline, e.g. sperm [46, 47]. Aligning long reads to this pangenome could detect the large number of recombination events present in sperm, similar to the path congruence analysis presented here, noting where reads change onto a different haplotype path. In the intermediate future, cattle tend to have highly related pedigrees due to the extensive use of breeding bulls, and so reanalysis of existing male long read cohorts with known pedigree (e.g., [48]) may already provide evidence of SV-recombination interactions.

Methylation, specifically 5mC, is also becoming increasingly accessible directly from long read sequencing. We find compelling evidence for several instances of allele-specific methylation or joining phaseblocks in the absence of SNPs. However, we found the initial differences in methylation signal are dominated by genomic mutations disrupting CpG motifs, rather than epigenetic-only change, requiring careful comparison across samples. Furthermore, the presence of intermediate methylation calls (i.e. between 30% and 70% methylated) further confounds analyses. However, improvements in sequencing chemistry and basecalling have already increased accuracy of 5mC methylation calls and enable the use of other methylation markers, which may provide additional evidence of haplotype-of-origin to detect recombination events.

## Conclusions

While massive pedigrees and genotype arrays are useful for confidently identifying recombination hotspots, we show related haplotype-resolved assemblies and pangenomes can accurately identify recombination events with high resolution. This approach also can identify haplotype status in the absence of heterozygous SNPs by incorporating structural variants and methylation signals, further improving the resolution of recombination maps. However, these approaches will benefit from further accuracy improvements for both assembly base-level quality and single CpG methylation scores to minimise false positive recombination events.

## List of abbreviations

ASM: Allele-specific methylation
CpG: Cytosine-phosphate-guanine
DMS: Differentially methylated sites
IBD: Identity by descent
QV: Quality value (in context of base-level accuracy)
RoH: Run of homozygosity
SNP: Single nucleotide polymorphism
SV: Structural variant
UPGMA: Unweighted pair group method with arithmetic mean

## Ethics approval and consent to participate

The sampling of blood was approved by the veterinary office of the Canton of Zurich (animal experimentation permit ZH137/2023).

## Consent for publication

Not applicable

## Availability of data and materials

Sequencing data for four Simmental genomes are public (PRJEB42335, PRJEB72196, and PRJNA677947), and the short sequencing reads of the Simmental dam of the Eringer x Simmental F1 (SAMEA115121771). New short sequencing reads for the Eringer sire of the F1 and the F1 itself have been made public (SAMEA121233886 and SAMEA121233887 respectively at study accession PRJEB28191), as well as the long sequencing of the F1 (SAMEA121647171 at study accession PRJEB42335). All workflows related to these analyses are available at https://github.com/ASLeonard/simbling.

## Authors’ contributions

ASL assembled the non-public genomes, constructed the pangenomes, and conducted all analyses. ASL and HP wrote the manuscript.

## Acknowledgements

We thank Cord Drögemüller for providing access to the SWISS_1 Simmental genome assembly. We also thank Xena Mapel for extracting the F1 and sire DNA for the Eringer x Simmental cross and Naveen Kadri for useful discussions regarding expected meiotic crossovers in half-siblings.

