## Supplementary material for "Identifying crossovers in a cattle pangenome containing haplotype-resolved assemblies from half-siblings": supplementary_material.docx

Supplementary Figure 1 (External PDF). Jaccard distance binned every 100 Kb across all pairs of assemblies for all chromosomes. Long stretches at zero indicate near-identical paths, with putative recombination events occurring at rapid increases of distance.

Supplementary Figure 2 (External PDF). Heterozygous maternal SNPs binned every 100 Kb where the two half-sibling haplotype assemblies had the same (blue) or opposite (orange) genotype across all chromosomes. Black circles mark every bin with nonzero variants, such that large gaps without markers indicate extended RoHs (e.g. chromosomes 3, 7, and 17).

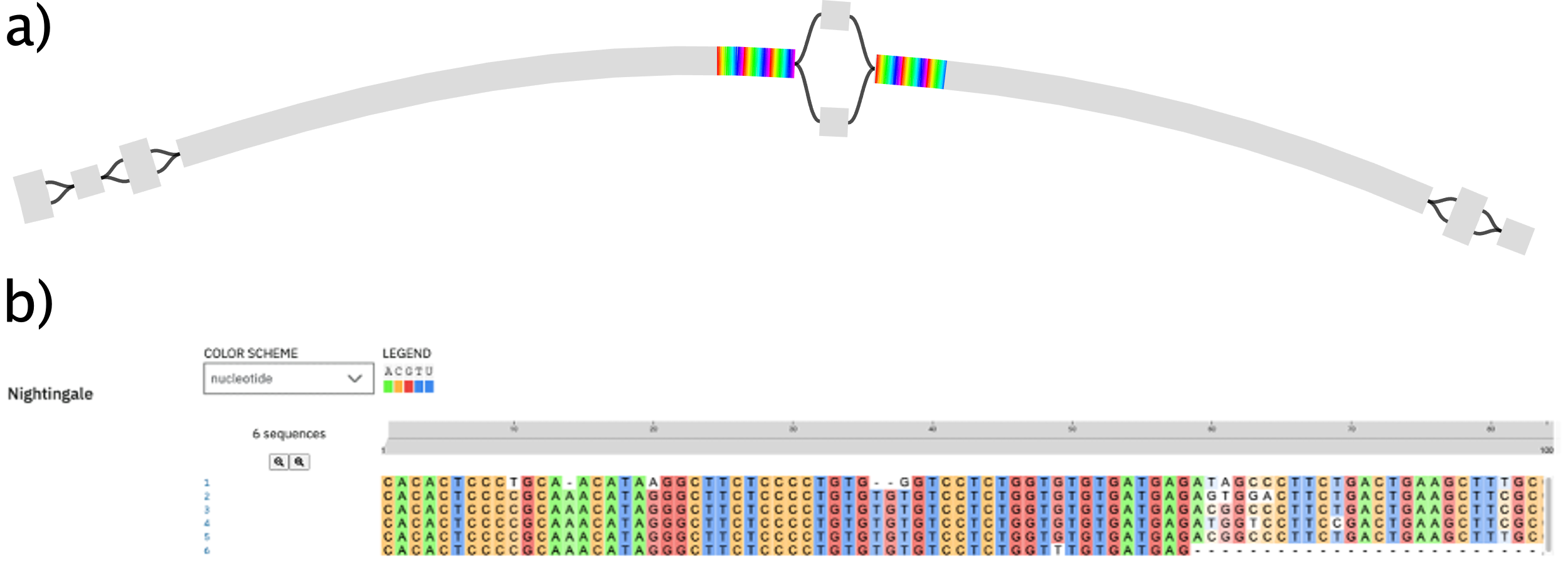

Supplementary Figure 3. (a) Nucleotide sequence of the PRDM9 motif mapped into the Simmental pangenome graph. In this region, there is only a small bubble corresponding to one SNP. Each full rainbow colouring (red to violet) corresponds to a full PRDM9 motif, while a truncated rainbow corresponds to a truncated motif. (b) USA_1 had the structure 1>2>2>3>2>4>6. All Swiss assemblies had 1>2>2>5>2>4>6, where motifs 3 and 5 differ by a single base in the 81^st^ position (C-to-T).

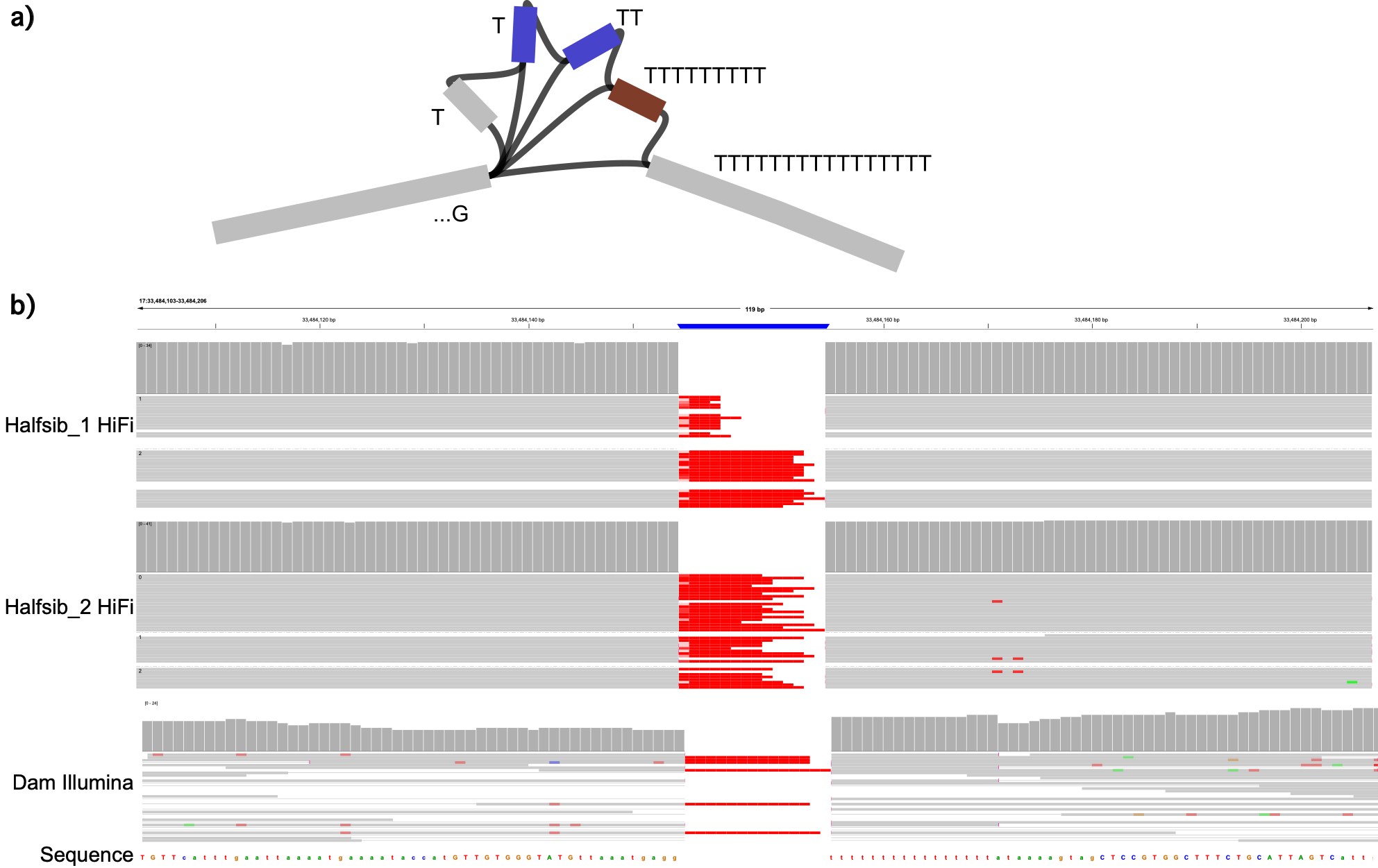

Supplementary Figure 4. (a) A long homopolymer stretch of T’s leads to many alleles in the graph, with each path having a unique allele. Both half-sibling assemblies take the brown node, while the assembly from half-sibling 1 also takes the two blue nodes. (b) HiFi reads from both half-sibling F1s have limited consensus on homopolymer length, leading to artefactual differences in the assemblies leading to spurious heterozygous alleles. Maternal short read sequencing is more robust to homopolymer errors and supports a more consistent consensus length.

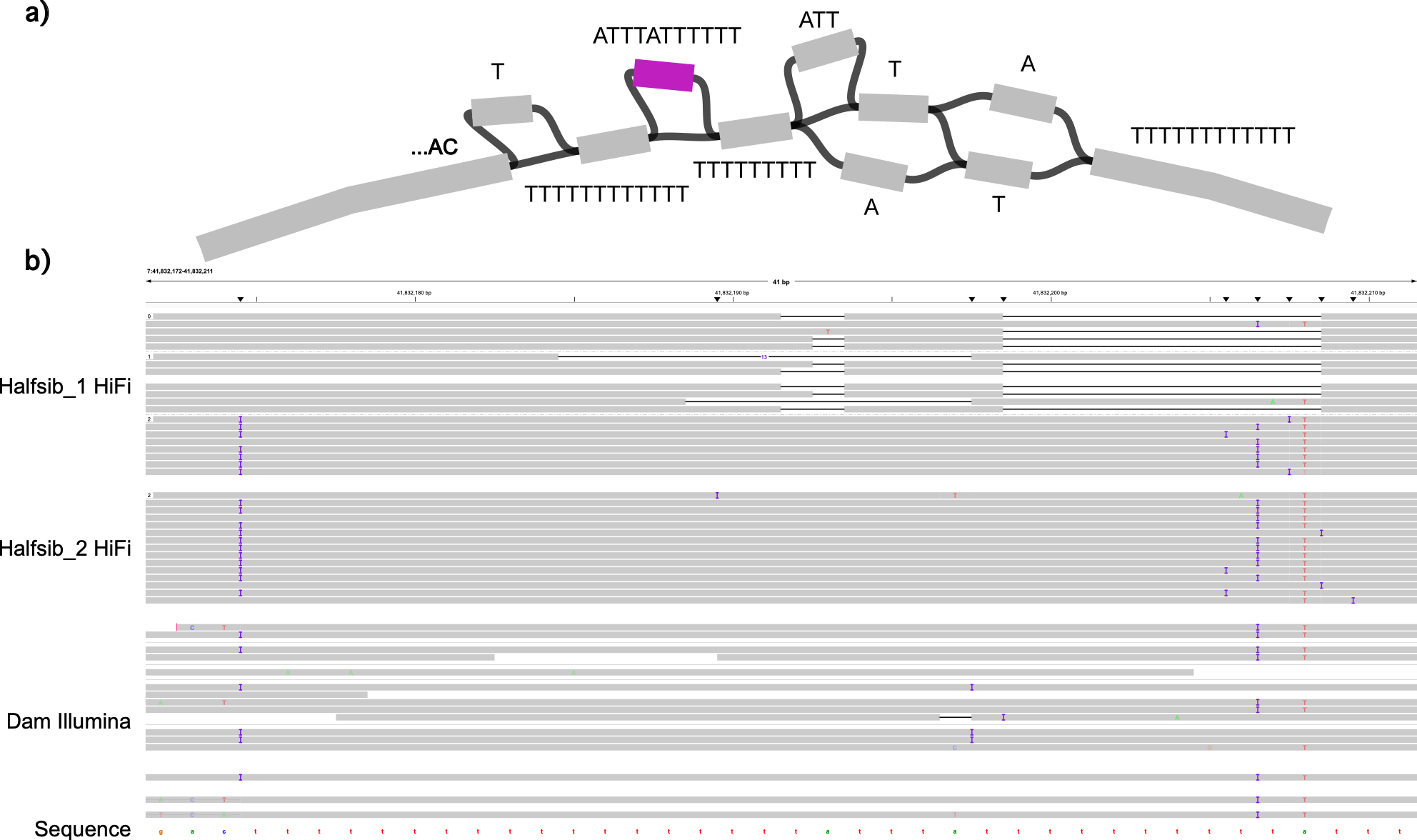

Supplementary Figure 5. (a) The Halfsib_1 path does not take the indicated node, equivalent to an 11 bp deletion relative to the USA_1 reference and other assemblies. (b) The 11 bp deletion appears in reads triobinned into the paternal Gyr haplotype of Halfsib_1 or ambiguous haplotype, indicating it is not present in the maternal Simmental haplotype. However, potentially due to poor graph phasing during assembly due to the repetitive nature of the region, the 11 bp deletion still erroneously ends up in the Simmental haplotype assembly, leading to a spurious heterozygous allele in the graph.

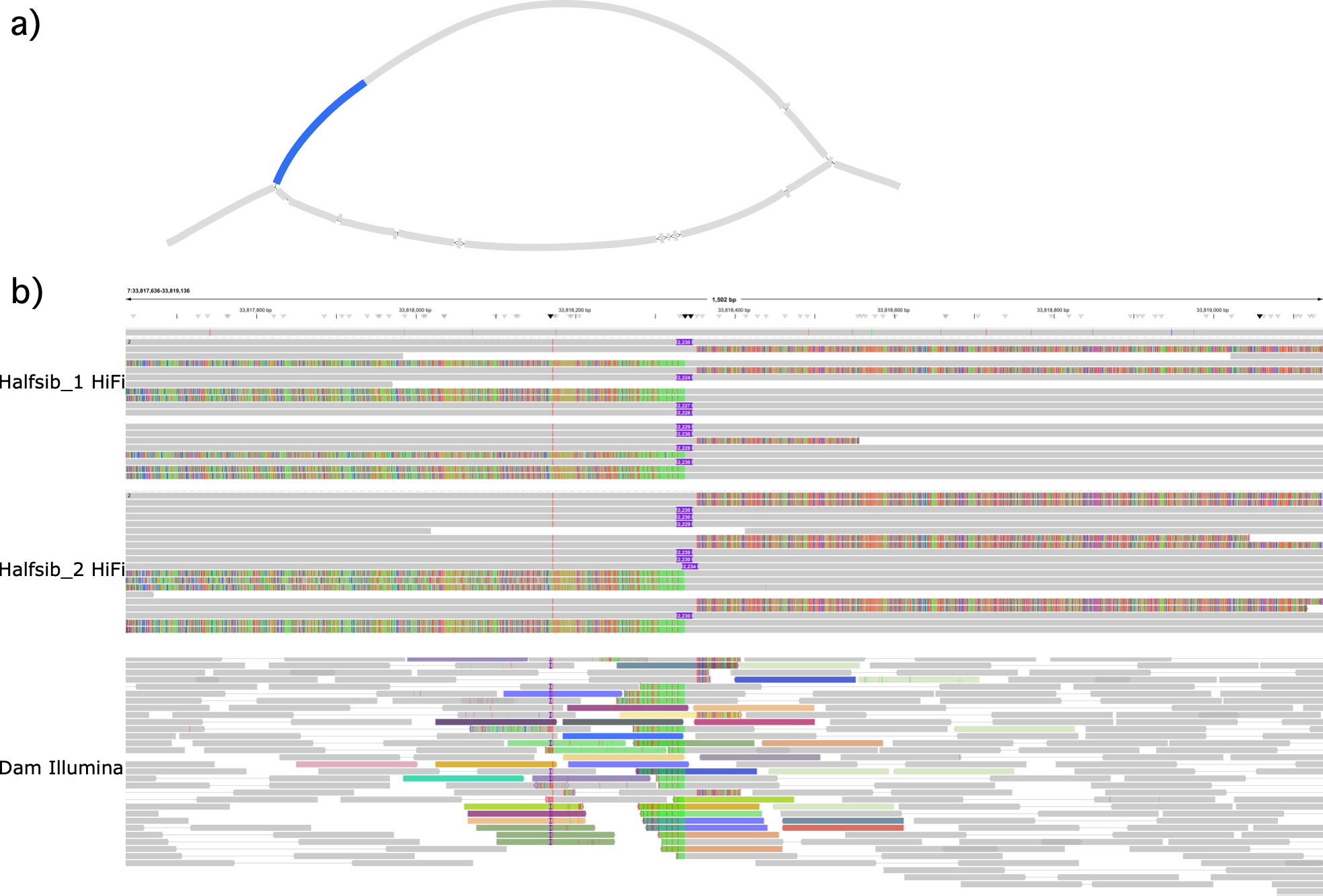

Supplementary Figure 6. (a) Incorrect pangenome deconstruction into VCF can lead to misleading heterozygous variants. Both half-sibling assemblies take the upper allele path (where the 2.3 Kb insertion is marked in blue), which shares some sequence with the lower allele path. After processing with vcfwave, Halfsib_2 has the 2.3 Kb insertion while Halfsib_1 is erroneously called as reference. More recent versions of vcfwave (v1.0.13+) appear to correct this issue. (b) The HiFi read alignments from both half-sibling F1s (and the maternal short read sequencing) support the 2.3 Kb insertion.

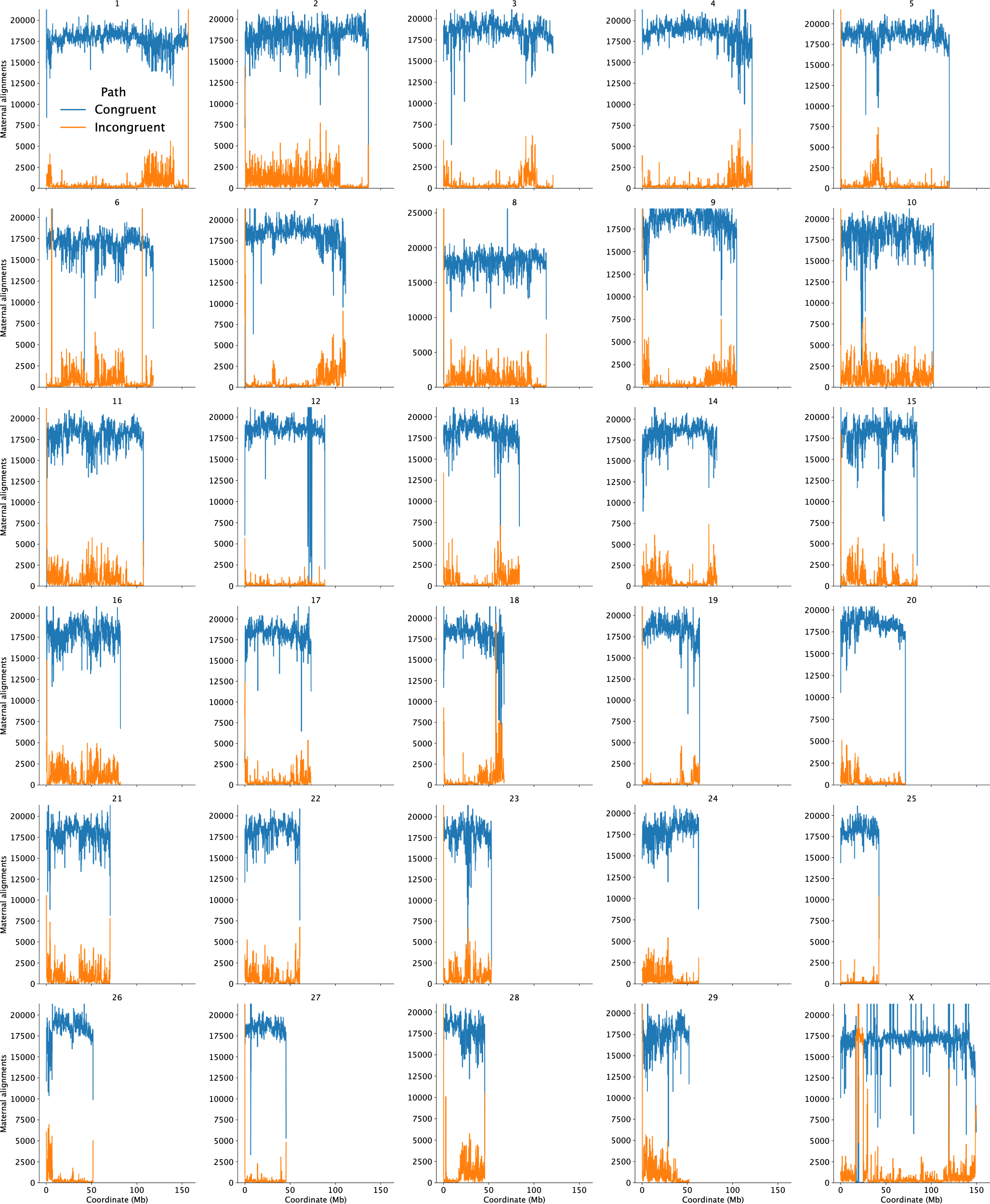

Supplementary Figure 7. Maternal short read sequencing alignment to the pangenome graph across all chromosomes. Alignment paths that match either haplotype walk are congruent (blue), indicating the half-sibling assemblies likely represent both maternal alleles. Incongruent (orange) paths suggest the maternal allele is not represented in the graph, i.e. both half-siblings inherited the same maternal allele at a heterozygous locus.

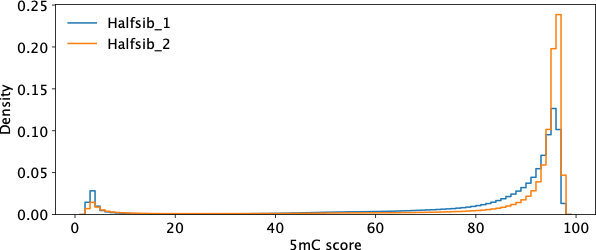

Supplementary Figure 8. Distribution of piled up 5mC scores from pb-cpg-tools. Peaks are largely bimodal and dominated by methylated CpGs, although Halfsib_1 has a more diffuse peak leading to ambiguous/intermediate methylation calls.

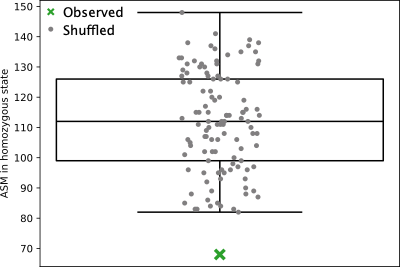

Supplementary Figure 9. Overlap of allele-specific methylation and “in-phase” inheritance (green X) and randomly permuted equal intervals (grey circles). Fisher’s exact test of the number of overlaps compared to the naïve expectation (68 versus 112 out of 207 total) was statistically significant (one-sided p=9.4×10^-6^), while the z-score of the observed arrangement compared to 100 random permutations was also statistically significant at p=0.0039).

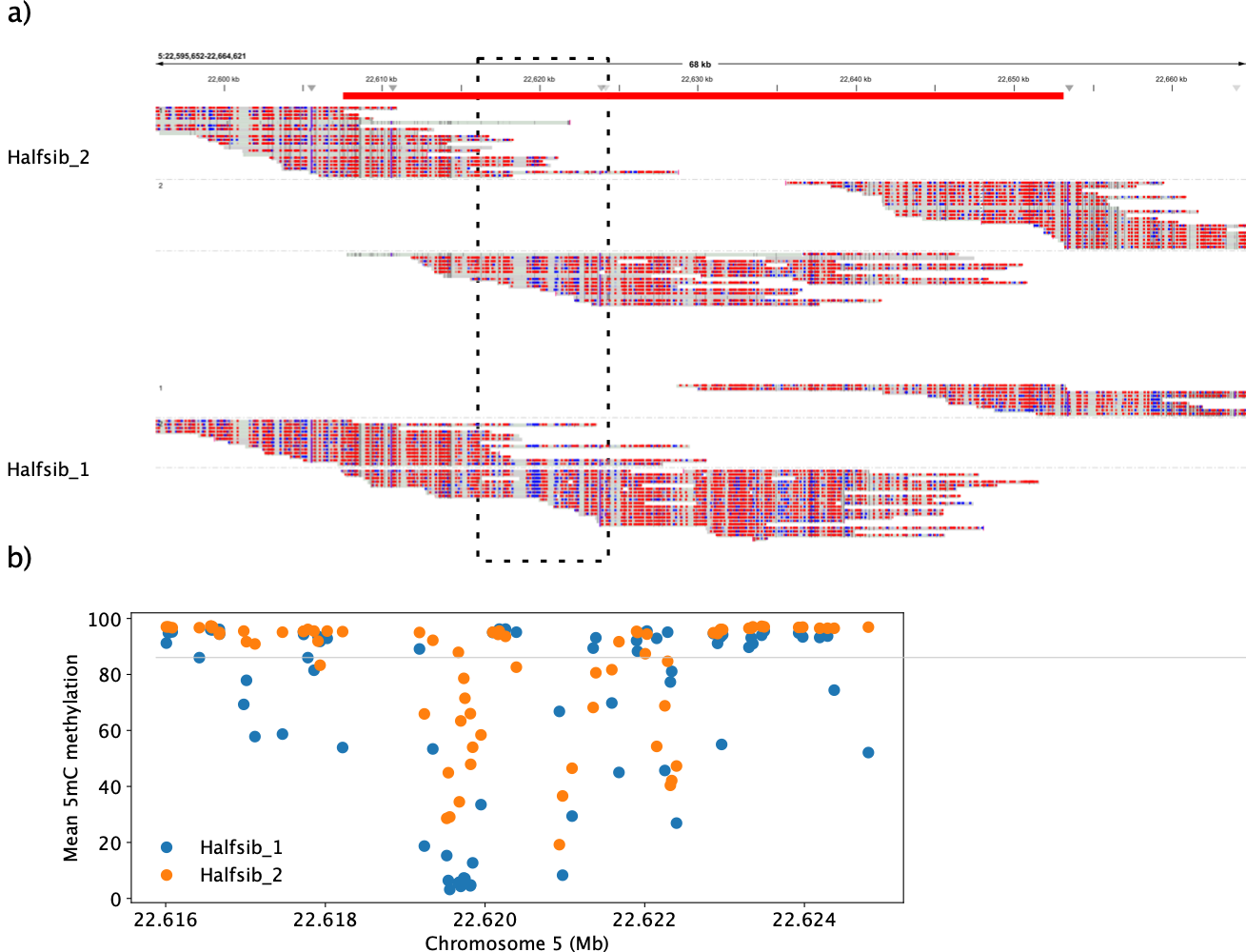

Supplementary Figure 10. (a) IGV screenshot of the SNP-based phasing of long reads, where reads are separated into their known haplotype-of-origin and then further divided into tagged phase. Differential 5mC methylation is shown as blue (hypomethylated) and red (hypermethylated). The red bar at the top indicates the extent of the unresolved phase block before considering the methylation signal. The black box indicates the region containing the most informative meth-mers used to phase reads. (b). Mean 5mC methylation for the two half-sibling haplotypes over the boxed region in (a) which was informative for methylation-based joining of phaseblocks.

**Supplementary Tables**

*Supplementary Table 1. Simmental assemblies used in the pangenome. Assembly metrics for the new Simmental haplotype (Halfsib_2) are reported for haplotype phased N50, merqury QV, and compleasm gene completeness.*

*Supplementary Table 2. Recombination events estimated per chromosome from both Jaccard path similarity and maternal-SNP phasing between the two half-siblings and the each of the half-sibling-cousin pairings.*

*Supplementary Table 3. Extended runs of homozygosity in dam as identified through biallelic SNPs called from short read sequencing. Although heterozygous SNPs are incredibly depleted (as expected in RoHs), there are still many heterozygous indels due to various homopolymer/short tandem repeat errors.*

*Supplementary Table 4. Differentially methylated regions (DMR) in the half-sibling HiFi reads, indicating the number of differentially methylated CpGs (>70% difference) and the approximate span of the region. Case indicates whether the DMR falls within a region that is expected to be in phase or antiphase, or the transition of phase across the RoH. The single DMR on chromosome 7 is a putative error since the RoH is expected to still be the same inherited haplotype, although this is the smallest and weakest DMR we identified.*

*Supplementary Table 5. Phaseblock joins possible with methylation signals (“meth-mers”) from pomfret, improving existing SNP-based phasing of HiFi reads. Cis- and trans-joins refer to reconciling arbitrary phase across phaseblocks (e.g. joining HP:i:0 tagged reads to HP:i:0 [cis] or HP:i:1 [trans]), and not proximal versus distal joins.*
