## Supplementary figures and images for "Identifying crossovers in a cattle pangenome containing haplotype-resolved assemblies from half-siblings"

### Supplementary_Figure_1.pdf

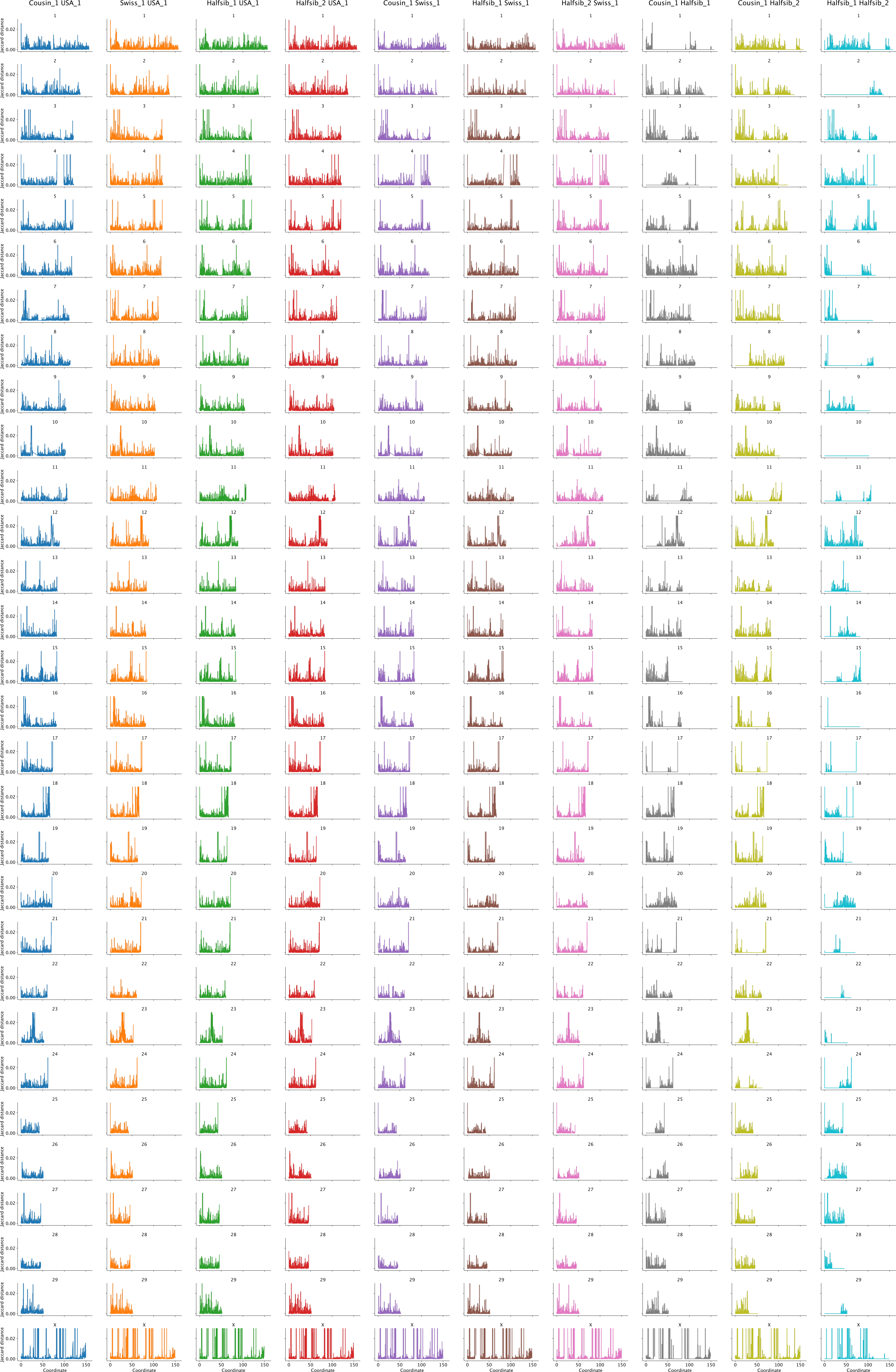

### Supplementary_Figure_2.pdf

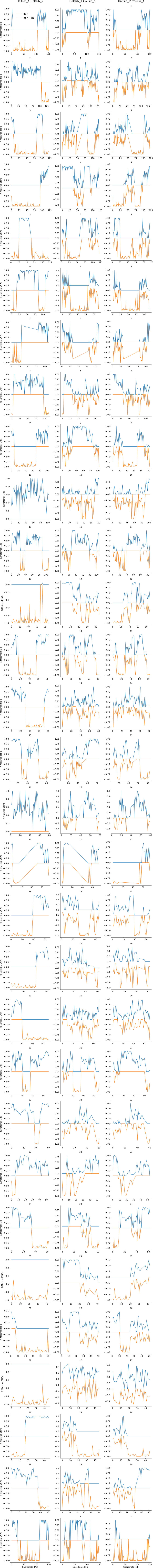
